# The scaffolding protein Cnk Interacts with Alk to Promote Visceral Founder Cell Specification in *Drosophila*

**DOI:** 10.1101/099986

**Authors:** Georg Wolfstetter, Kathrin Pfeifer, Jesper Ruben van Dijk, Fredrik Hugosson, Xiangyi Lu, Ruth Helen Palmer

## Abstract

In *Drosophila*, the receptor tyrosine kinase Alk and its ligand Jeb are required to drive founder cell (FC) specification in the visceral mesoderm (VM). Alk-signalling activates downstream MAPK/ERK- and PI3K-pathways in human and *Drosophila* but little is known about immediate downstream signalling events. Here we report that the scaffolding protein Cnk interacts directly with Alk via a novel c-terminal binding motif. Cnk is required for Alk-signalling as ectopic expression of the minimal interaction motif as well as loss of maternal and zygotic *cnk* blocks visceral FC-formation, resembling the phenotype of *jeb* and *Alk* mutants. We also show that the Cnk-interactor Aveugle/Hyphen (Ave/HYP) is critical, while the (pseudo-) kinase Ksr is not required for Alk-signalling in the developing VM. Taken together, Cnk and Ave represent the first molecules downstream of Alk whose loss genocopies the lack of visceral FC-specification of *Alk* and *jeb* mutants indicating an essential role in Alk-signalling.

## INTRODUCTION

Receptor tyrosine kinase (RTK)-signalling plays an essential role in development by transducing external signals into the nucleus and other cellular compartments thereby altering gene expression and promoting intracellular changes. The hallmarks of RTK-signalling are highly conserved amongst eukaryotic organisms and involve ligand-dependent activation of a transmembrane receptor protein-tyrosine kinase and the recruitment of a canonical series of intracellular signalling modules and cascades. The mitogen-activated protein kinase/extracellular-signal regulated kinase (MAPK/ERK) pathway is a prominent example of such a signalling cascade employing the Ras-GTPase cycle and the serine/threonine kinases Raf (MAPKKK), MEK (MAPKK), and MAPK/ERK to transduce signals downstream of an activated receptor (Widmann et al. 1999) (Kolch 2000) (Schlessinger 2014).

In addition to the core components involved in MAPK/ERK-signalling, many other factors have been identified over recent decades that contribute to or modulate the overall pathway activity. One such factor is the kinase suppressor of Ras (Ksr) that was identified by mutagenesis-screens in Ras-sensitised genetic backgrounds in both *Drosophila melanogaster* and *Caenorhabditis elegans* (Kornfeld et al. 1995) (Sundaram and Han 1995) (Therrien et al. 1995). Ksr is an evolutionary conserved protein kinase, closely related to Raf that operates downstream of a variety of RTKs such as FGFR, Torso, Sevenless, and EGFR (Kornfeld et al. 1995) (Sundaram and Han 1995) (Therrien et al. 1995) (Cabernard and Affolter 2005). However, due to inconsistent findings regarding the catalytic activity of its kinase domain, the role of Ksr has remained controversial. Different models proposed different roles for Ksr as a Raf-activator in parallel or downstream of Ras, as an activating kinase in a different signalling branch or as a scaffolding protein for the assembly of Raf/MEK protein complexes (Kornfeld et al. 1995) (Sundaram and Han 1995) (Therrien et al. 1995) (Morrison 2001) (Roy et al. 2002). There is no evidence for a direct interaction between Ksr and Ras (Claperon and Therrien 2007). However, recent findings have revealed that side-to-side dimerization between Ksr and Raf can invoke intrinsic Raf activation (Rajakulendran et al. 2009) providing a mechanism for Ksr in RTK-signalling that does not require its functional kinase domain.

A genetic modifier screen employing ectopic expression of a dominant-negative, chimeric version of Ksr (Therrien et al. 1996) in the fly eye led to the identification of another critical factor for Ras/ERK-signalling named <u>C</u>onnector e<u>n</u>hancer of <u>k</u>inase suppressor of Ras (Cnk) (Therrien et al. 1998). The *cnk* locus encodes a large protein of 1,557 amino acids containing an n-terminal <u>S</u>terile <u>A</u>lpha <u>M</u>otif (SAM), followed by a <u>C</u>onserved <u>R</u>egion <u>I</u>n <u>C</u>nk (CRIC), a PDZ domain, Proline-rich motifs and a Pleckstrin Homology (PH) domain. The protein structure suggests that Cnk performs versatile functions as a multi-domain protein scaffold. Like Ksr, Cnk functions downstream of various RTK-signalling events including EGFR-mediated patterning of wing disc territories and FGFR/EGFR-dependent air sac development in the dorsal thorax (Baonza et al. 2000) (Cabernard and Affolter 2005). Interestingly, while ectopic expression of the Cnk n-terminal region enhances the effects of activated Ras^V12^ independently of MAPK/ERK activation in the *Drosophila* eye, the c-terminal region binds to and regulates Raf by a bi-modal mechanism involving a Raf inhibitory region (RIR) and a phosphorylation site for the Src family kinase Src42A (Therrien et al. 1998) (Therrien et al. 1999) (Douziech et al. 2003) (Laberge et al. 2005). Thus, Cnk has complex roles functioning as a molecular scaffold to support Ksr-mediated Raf activation as well as to recruit and integrate additional signalling components such as Src42A.

During embryonic development in *Drosophila melanogaster*, the visceral mesoderm (VM) gives rise to a lattice of midgut muscles that ensheaths the larval midgut. The VM consists of initially undifferentiated myoblasts that become specified as either founder cells (FCs) or fusion competent myoblasts (FCMs). Subsequently, the FCs fuse one-to-one with the FCMs and eventually form the bi-nucleate visceral myotubes (Poulson 1950) (Campos-Ortega and Hartenstein 1985) (Martin et al. 2001) (Klapper et al. 2002) (Lee et al. 2006).

Earlier work employing phospho-specific antibodies to detect ERK-activation in the *Drosophila* embryo suggested a novel signalling pathway participating in the specification of VM cells (Gabay et al. 1997). The identity of the RTK involved was revealed by the discovery of a *Drosophila* orthologue of the Anaplastic Lymphoma Kinase (ALK) receptor, initially identified as part of a chimeric protein created by the 2;5 (p23:q35) translocation in human anaplastic large cell lymphoma cell lines (Morris et al. 1994) (Loren et al. 2001). In *Drosophila, Alk* is expressed in the segmental clusters that segregate from the dorsal trunk mesoderm to form the VM (Loren et al. 2001). The Alk protein can be detected at the membrane of all VM cells but only the distal arch within each cluster comes into direct contact with a secreted, small LDL-domain ligand named Jelly belly (Weiss et al. 2001) (Loren et al. 2001) (Englund et al. 2003) (Lee et al. 2003). Direct binding of Jeb to the extracellular part of Alk leads to the activation of a downstream signalling cascade eventually resulting in ERK phosphorylation (Englund et al. 2003) (Lee et al. 2003). As a consequence of this signalling event, expression of a FC-specific subset of genes including *Hand*, the *optomotor-blind-related-gene-1* (*org-1*) and *dumbfounded/kin of irreC* (*duf/kirre*) is initiated and/or maintained in these cells (Englund et al. 2003) (Lee et al. 2003) (Stute et al. 2004) (Varshney and Palmer 2006) (Schaub et al. 2012) (Schaub and Frasch 2013). Subsequent studies of *jeb* and *Alk* mutant embryos showed that Jeb/Alk-signalling is indeed crucial for visceral myoblasts to commit to the FC-fate. In the absence of either ligand or receptor, neither ERK-phosphorylation nor the expression of FC-specific marker genes in the VM is observed. Moreover, the visceral cells fail to undergo myoblast fusion and the VM subsequently disintegrates in *jeb* and *Alk* mutant embryos (Loren et al. 2001) (Loren et al. 2003) (Englund et al. 2003) (Lee et al. 2003) (Stute et al. 2004).

In addition to its role in visceral FC-specification, Alk-signalling is employed in a variety of other developmental processes. These include a larval brain-sparing mechanism in which tissue-autonomous Alk-signalling modifies Insulin/PI3-Kinase/Target Of Rapamycin (TOR) pathway activity to allow growth of CNS progenitors under nutrient restriction conditions (Cheng et al. 2011). Studies have also revealed roles for Alk-signalling during neuronal circuit formation in the fly visual system and in synaptic signalling at the larval neuromuscular junction (Bazigou et al. 2007) (Rohrbough et al. 2013) (Pecot et al. 2014). Moreover, Alk was found to act as an upstream activator of Neurofibromin-1 (NF1) linking its function to associated learning, memory formation and body size regulation (Gouzi et al. 2011). Despite our growing knowledge about the multiple roles of Alk-signalling and the variety of processes it is involved in, there is a gap in our present knowledge concerning the identity of the immediate downstream factors that transduce and distribute the initial activation signal towards the respective effectors.

In this study we identify the multi-domain scaffolding protein Cnk as a novel Alk binding partner and essential component in the Alk-signalling pathway. Cnk binds to the intracellular part of Alk via a c-terminal interaction motif. Loss of *cnk* function or expression of dominant-negative *cnk*-constructs in *Drosophila* interferes with Alk-signalling in multiple developmental contexts. Moreover, germline clone-derived embryos lacking maternal and zygotic Cnk fail to specify visceral FCs and do not develop a functional midgut. In agreement with its proposed function, epistasis experiments reveal that Cnk operates between Ras and Raf in the Alk-signalling pathway. Further targeted deletion of the minimal <u>A</u>lk <u>i</u>nteraction <u>m</u>otif (AIM) in Cnk results in a specific decrease of Jeb/Alk-induced ERK-phosphorylation within the visceral FC row. Interestingly, while the SAM-domain containing Cnk-binding partner Aveugle/Hyphen (Ave/HYP) is also essential for Alk-signalling, we find that Ksr is not required to drive Alk-signalling in the developing VM. Taken together, we identify a novel interaction motif that mediates binding of Cnk to Alk and show that both Cnk and its binding partner Ave are critical components of Alk-signalling in the developing *Drosophila* embryonic visceral mesoderm.

## RESULTS

### Cnk is a direct binding partner of the Alk receptor

In order to identify novel components of the Alk-signalling pathway in *Drosophila*, yeast two-hybrid (Y2H) screening was performed employing the intracellular portion (spanning amino acids Tyr1128 to Cys1701) of the *Drosophila* Alk protein, (subsequently referred to as Alk^ICD^) as bait. The screen was conducted with prey constructs derived from a random-primed cDNA-library extracted from adult fly heads. Thirty-six clones encoding portions of Cnk were identified as Alk^ICD^-interactors (Fig. 1A). Further analysis defined a minimal overlapping region of 42 amino acids that was sufficient to bind Alk (Ala1384-Ser1425, further referred to as the Cnk <u>A</u>lk <u>i</u>nteraction <u>m</u>otif or Cnk^AIM^) located in the functionally unannotated c-terminal region of Cnk (Fig. 1B). Thus, the Cnk^AIM^ region contains the minimal requirements to mediate the interaction of Cnk with the Alk^ICD^.

**Figure 1.**
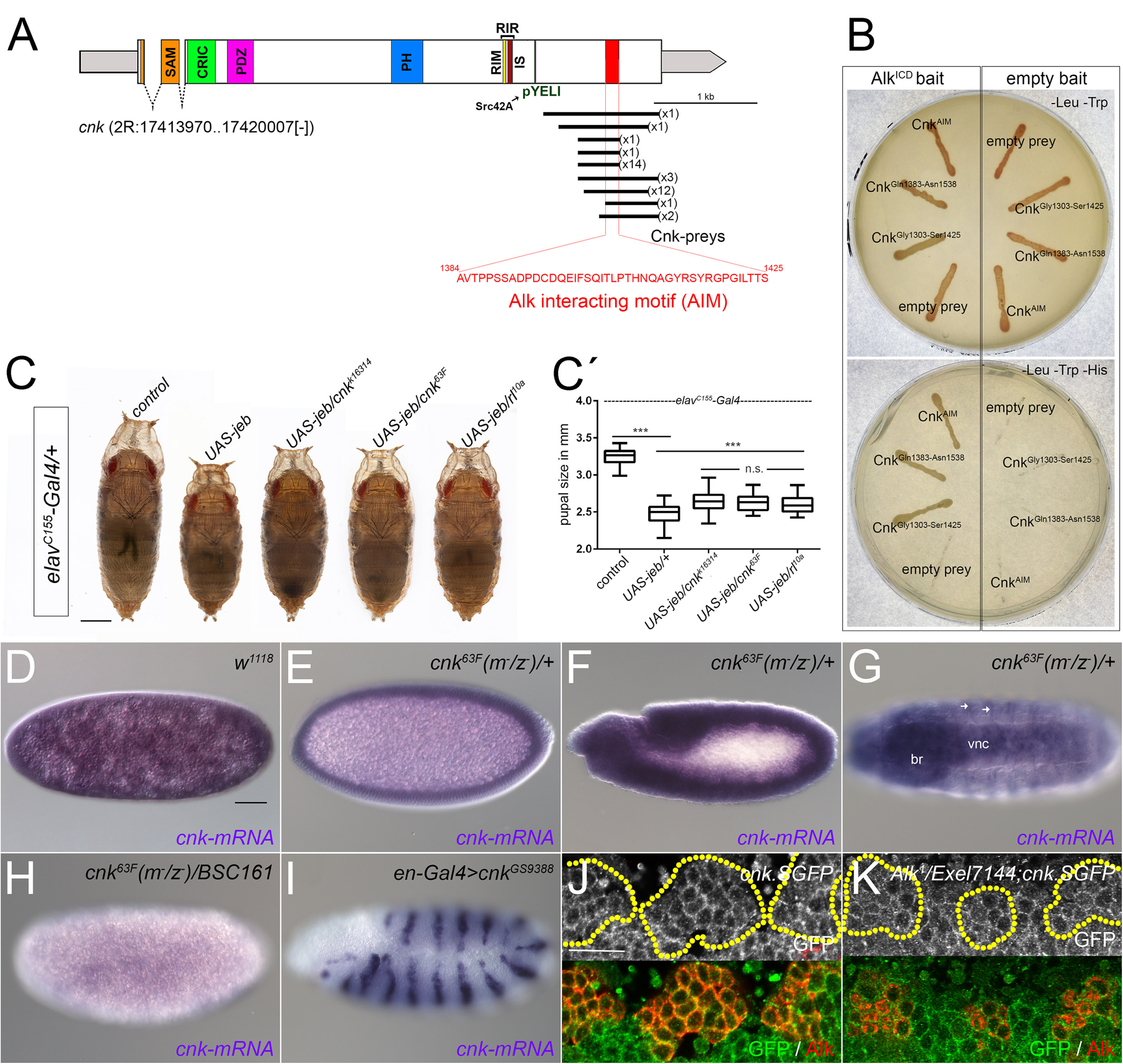
Alk and Cnk interact via a novel Alk interacting motif (AIM) **(A)** Schematic representation of the *cnk* locus (adapted and modified from Clapéron and Therrien (Claperon and Therrien 2007)) depicting the number and position of Alk^ICD^-interacting Cnk-preys obtained by Y2H analysis. Regions encoding for Cnk domains (SAM=sterile alpha motif, CRIC=conserved region in Cnk, PDZ=post synaptic density protein (PSD95), *Drosophila* discs large tumor suppressor (Dlg1), and zonula occludens-1 protein (zo-1), PH=pleckstrin homology domain, RIR=Raf inhibitory region, RIM=Raf interacting motif, IS=inhibitory sequence, pYELI=phosphorylation site for Src42A) are highlighted in different colours. UTRs appear narrower and in grey, introns as dashed lines. The position of the minimal <u>A</u>lk <u>i</u>nteracting <u>m</u>otif (AIM; Ala1384-Ser1425) is shown in red. **(B)** Yeast growth on double-selective transformation/mating-control (-Leu/-Trp) and triple selective (-Leu/-Trp/-His) media plates. Each streak corresponds to a diploid yeast colony co-expressing Gal4_BD-Alk^ICD^ (left half) or Gal4_BD (empty vector control, right half) and Gal4_AD-fusion proteins with the respective Cnk-prey fragments indicated. Alk^ICD^ specifically binds to Cnk fragments covering the region identified (A) and to the minimal overlapping region (Cnk^AIM^). **(C)** Cnk modifies *elav^C155^-Gal4>UAS-Jeb* induced pupal size reduction. Late stage female *Drosophila* pupae expressing *UAS-Jeb* pan-neuronally with *elav^C155^-Gal4.* Alterations in the genetic background are depicted over the respective pupa. **(C´)** Quantification of the Cnk-dependent modification of *elav^C155^-Gal4>UAS-Jeb* induced pupal size reduction. Variance analysis (one way ANOVA) indicated that the effects of genotypes was significant (F(_4_,_221_)=174,4, p<0.001, n=226) and pairwise Student’s t-test was employed to reveal the significance of the measured size-differences (***=p<0,001 or n.s.= not significant). **(D-I)** Whole mount *in situ* hybridisation on *Drosophila* embryos with a cnk-specific probe: Maternal *cnk* transcripts in a stage 2 *white^1118^* embryo (D), ubiquitous zygotic *cnk* expression in cnk^63F^(m^−^/z^−^)/+ embryos at stages 5 (E) and 10 (F). Tissue-specific expression of *cnk* is revealed in the brain (br), ventral nerve chord (vnc), and the peripheral nervous system (arrows) at the end of embryogenesis (G). Absence of cnk-transcripts in *cnk^63F^(m^−^/z^−^)/BSC161* embryos (H) and *en2.4-Gal4-driven* cnk-misexpression employing the *P{GSV6}GS9388* insertion (I) reveals specificity of the *cnk in situ* probe. **(J-K)** Cnk and Alk protein localisation in stage 10/11 control (J) and transheterozygous *Alk^1^/Exel7144* (K) embryos revealed by the FlyFos TransgeneOme library line *JTRG1248 (Cnk.SGFP).* The majority of Cnk.SGFP was observed in the vicinity of the plasma membrane in the presence and absence of functional Alk protein. Scale bars: 500 μm in C, 50 μm in D, 10 μm in J.

### Cnk modulates Jeb/Alk-signalling in the CNS

To verify the *in vivo* relevance of the Alk-Cnk interaction, we first tested the ability of Cnk to modify the previously described pupal size reduction phenotype caused by elevated Alk-signalling (Gouzi et al. 2011). The *Drosophila* Alk-ligand Jelly belly (Jeb) was expressed with the pan-neuronal *elav^C155^-Gal4* driver resulting in a pupal size reduction of about 20% when compared to *elav^C155^-Gal4/+* controls (Fig. 1C). Introducing single copies of the loss of function alleles *cnk^k16314^* or *cnk^63F^* (see Material and Methods and Fig. 3-figure supplement 1 for further information regarding *cnk* alleles) in the Jeb-overexpression background resulted in a significant (***= p<0.001 in pairwise two-tailed Student’s t-test; n>45) increase in pupal size, indicating that loss of Cnk counteracts the effect of enhanced Jeb/Alk-signalling (Fig. 1C, quantified in C´). As control we reduced the dose of the *Drosophila* ERK1/2 homologue *rolled*, employing the chromosomal deletion *Df(2R)rl^10a^*, which resulted in a comparable increase of pupal size as observed with reduction of Cnk. Taken together, these results supported a role for Cnk as novel downstream component of Jeb/Alk-signalling.

### Cnk is expressed in the VM but does not depend on Alk for its localisation

One of the most prominent functions for Alk-signalling in *Drosophila* is the specification of visceral muscle FCs in the developing embryo (Lee et al. 2003) (Englund et al. 2003) (Stute et al. 2004). Before analysing a possible role for Cnk in this process we investigated whether Cnk and Alk are co-expressed in the embryonic VM during Jeb/Alk-dependent FC-specification. First, we employed *cnk*-specific antisense probes in *in situ* hybridisations on *Drosophila* embryos (Fig. 1D-G, specificity controls in H-I). Our analysis revealed a strong maternal component for *cnk* suggesting that all cells in the embryo are supplied with Cnk (Fig. 1D). Zygotic *cnk* transcription in progeny of *cnk^63F^* germ line clone-producing females crossed to *w^1118^* control males was detectable in all cells at the syncytial blastoderm stage (Fig. 1E) and remained ubiquitously expressed during germ band elongation (Fig. 1F). After germ band retraction, ubiquitous expression had declined and *cnk* was specifically expressed in the brain (br), ventral nerve cord (vnc) and the peripheral nerves (arrows) at the end of embryogenesis (Fig. 1G). In addition to *in situ* hybridisation, we employed the multi-tagged FlyFos TransgeneOme (fTRG) library line *fTRG1248* (subsequently referred to as *cnk.SGFP*) that carries an extra copy of the *cnk* locus encoding a c-terminally GFP-tagged variant of Cnk expressed under the control of its endogenous regulatory elements (Sarov et al. 2016). Notably, *cnk.SGFP* was able to rescue lethality caused by *cnk* lof-mutations (Supplementary file 1, Fig. 1-figure supplement 1), indicating that Cnk.SGFP can functionally substitute for the wild-type protein. At stage 10/11 Cnk.SGFP could be detected in all cells of the ectoderm and the underlying mesoderm including Alk-positive cells (Fig. 1J). The majority of the protein was localised in close proximity to the plasma-membrane (Fig. 1J) and this localisation of Cnk.SGFP was maintained in *Alk^1^/Df(2r)Exel7144 (Exel7144*, aa Alk deficiency)embryos that express one copy of a truncated Alk protein lacking the Alk^ICD^ suggesting that Cnk-localisation does not depend on Alk (Fig. 1K).

### Ectopic expression of the AIM region of Cnk interferes with *Alk*-signalling

To further examine a possible function for the interaction between Alk and the Cnk^AIM^, we generated *UAS*-constructs encoding a tandem array of five copies of Cnk^AIM^, c-terminally tagged with 3xHA (*UAS-5xcnk^AIM^.3xHA*). Since our previous results indicated that Cnk is localised in the proximity of the plasma membrane, we used the n-terminal myristoylation sequence of *Src42A* to generate a membrane-associated variant (*UAS-Myr⸬5xcnk^AIM^.3xHA*). We first studied the effects of 5xCnk^AIM^ upon ectopic expression of Alk in the developing eye with the *sevEP-Gal4* driver (Fig. 2A-E). While expression of Alk alone (Fig. 2C) but not of 5xCnk^AIM^ (Fig. 2A, B) led to severe defects in ommatidia formation in the anterior part of the eye, the combined expression of Alk and 5xCnk^AIM^ substantially restored a “wild-type” eye morphology (Fig. 2D, E). In agreement with our assumption that the membrane localisation of Cnk might influence its function, the *Myr⸬5xcnk^AIM^.3xHA* construct supressed the effects of ectopic Alk-signalling (Fig. 2E) more effectively than *5xcnk^AM^.3xHA* (Fig. 2D). We next tested the ability of 5xCnk^AIM^ to interfere with endogenous Alk-signalling during embryonic VM-development. Therefore, we expressed 5xCnk^AIM^ with the mesodermal *P{GAL4-twi.2xPE}* driver (henceforth referred to as *2xPE-Gal4*) and employed *rP298-lacZ* or *HandC-GFP* as FC-reporters (Nose et al. 1998) (Sellin et al. 2006) in combination with antibody staining against Alk or Fasciclin III (FasIII) to reveal visceral mesoderm and muscle morphology (Fig. 2F-K). Ectopically expressed 5xCnk^AIM^.3xHA exhibited a cytoplasmic expression pattern (Fig. 2G), while the Myr⸬5xCnk^AIM^.3xHA construct was membrane localised (Fig. 2H). Expression of the *rP298-lacZ* FC-marker was severely reduced in the VM of *2xPE>5xcnk^AIM^.3xHA* and *2xPE>Myr⸬5xcnk^AIM^.3xHA* embryos when compared with controls (Fig. 2 F-H). Moreover, the chambered midgut surrounded by HandC-GFP- and FasIII-positive visceral myotubes (Fig. 2I) was not formed at the end of embryogenesis (Fig. 2J, K) upon ectopic expression of either 5xCnk^AIM^.3xHA or Myr⸬5xCnk^AIM^.3xHA. The presence of *rP298-lacZ* expression in somatic FCs (Fig. 2G, H arrowheads) and the body wall muscles (Fig. 2J, K asterisks) in late stage *2xPE>5xcnk^AIM^* embryos further indicated that ectopic expression of 5xCnk^AIM^ affected visceral FC-formation but not somatic muscle development suggesting a specific effect on Alk-signalling in the embryonic VM.

**Figure 2.**
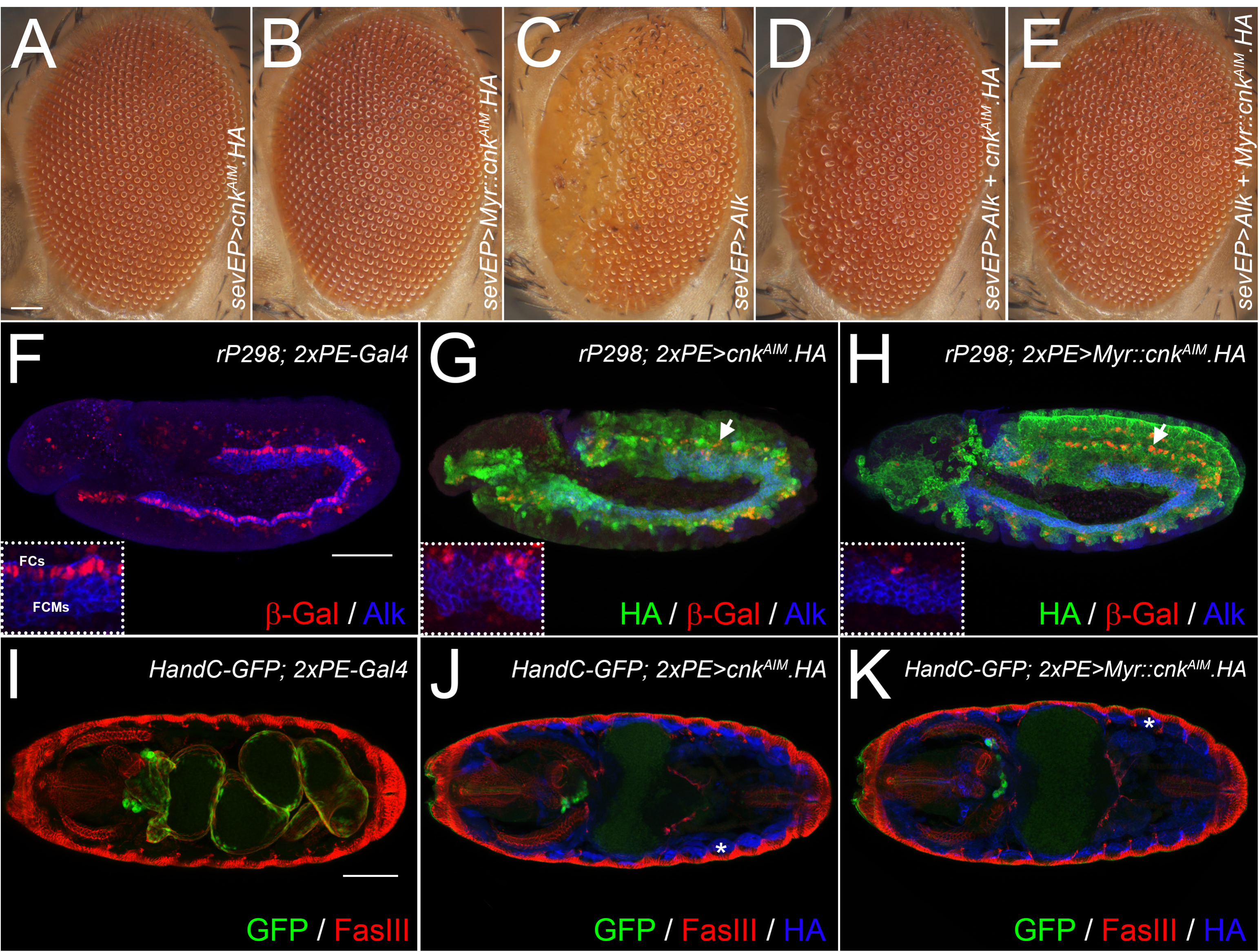
Ectopic expression of Cnk^AIM^ inhibits Alk-signalling. **(A-E)** Cnk^AIM^ inhibits ectopic Alk-signalling in the fly eye. Heads of adult female flies expressing the indicated *UAS*-transgene under the control of the *sev-Gal4* driver are depicted. Ectopic expression of HA-tagged 5xCnk^AIM^ (A), HA-tagged 5xMyr-Cnk^AIM^ (B) tandem arrays or *UAS-Alk* (C). Alk-induced defects can be rescued by co-expressing Alk with either HA-tagged 5xCnk^AIM^ (D) or HA-tagged 5xMyr-Cnk^AIM^ (E). **(F-H)** Cnk^AIM^ inhibits Alk-signalling in the embryonic VM. The trunk VM of stage 11/12 embryos was visualised by antibody staining against β-Galactosidase (β-Gal, red) to reveal *rP298-LacZ* reporter gene expression and anti-Alk (blue). Expression of either HA-tagged 5xCnk^AIM^ (G) or HA-tagged 5xMyr-Cnk^AIM^ (H) was detected by anti-HA-tag antibody staining (HA, green). *rP298-LacZ* expression observed in the visceral FC row (FCs, FCMs are fusion competent myoblasts) of control embryos (F) is strongly reduced by the ectopic expression of HA-tagged 5xCnk^AIM^ in the cytoplasm (G) or at the membrane (H). In contrast, *rP298-lacZ* expression in the somatic mesoderm (arrows in G & H) is unaffected. **(I-K)** Stage 16 embryos stained for *HandC-GFP* reporter gene expression (GFP, green), Fasciclin III (FasIII, red) and anti-HA (HA, blue). Control embryos (I) exhibit a four-chambered midgut surrounded by HandC-GFP- and FasIII-positive muscles. (J, K) Embryos expressing either HA-tagged 5xCnk^AIM^ (J) or HA-tagged 5xMyr-Cnk^AIM^ (K) under control of the *twist.2xPE* driver (*2xPE*) lack midgut muscles and do not form a midgut. The normal appearance of somatic muscles (asterisk in G, H revealed by HA-staining in blue) suggests a VM-specific role for Cnk^AIM^. Scale bars: 50 μm.

### Cnk function is required for visceral mesoderm development in the *Drosophila* embryo

The interaction between Cnk and the Alk^ICD^ in the Y2H assay as well as the ectopic expression of the Cnk^AIM^ led us to ask whether endogenous Cnk function might be critical for Alk-signalling in the *Drosophila* embryo. As *Alk, jeb* and *cnk* are located on the right arm of the second chromosome, we first confirmed that the *cnk* alleles used in this study complemented the loss of function mutations *Alk^1^* and *Alk^10^*, as well as *jeb^weli^, jeb^k05644^* and *Df(2R)BSC199*, a deficiency for *jeb* (Hugosson et al. 2014). Further complementation tests employing the *cnk* deficiency *Df(2R)BSC161* revealed that all analysed zygotic *cnk* mutants alleles appeared to be viable until late larval or pupal stages contradicting our assumption for a critical role of *cnk* in visceral FC specification. Moreover, analysis of VM-development in control siblings (Fig. 3A, D) and zygotic cnk-mutant embryos (Fig. 3B, E) employing *rP298*, and *HandC-GFP* as FC markers in combination with antibody staining against Alk or FasIII did not reveal a loss of FCs or later defects in the visceral muscle. Given the maternal *cnk* component revealed by our *in situ* analysis (Fig. 1D), we hypothesised that maternally derived Cnk could mask a potential visceral phenotype in zygotic *cnk* mutants, prompting us to examine germ line clone-derived embryos lacking maternal and zygotic (m^−^/z^−^) *cnk* (Fig. 3C, F). In agreement with an important role for Cnk in Jeb/Alk-signalling, we observed a VM-phenotype characterised by complete loss of FCs at stage 11/12 (Fig. 3C) and absence of differentiated midgut muscles at the end of embryogenesis (Fig. 3F). This fully penetrant phenotype was observed in maternal and zygotic (m^−^/z^−^) embryos of all *cnk* alleles analysed (Fig. 3-figure supplement 1) and was indistinguishable from the characteristic visceral defects of *jeb* and *Alk* mutant embryos (Fig. 3G, H, J and K) (Englund et al. 2003) (Lee et al. 2003) (Stute et al. 2004). Finally, Cnk.SGFP was able to restore visceral FC formation and the development of midgut muscles in maternal and zygotic (m^−^/z^−^) *cnk* embryos (Fig. 3I, L), providing further evidence that *cnk* specifically functions in the Jeb/Alk-signalling pathway.

**Figure 3.**
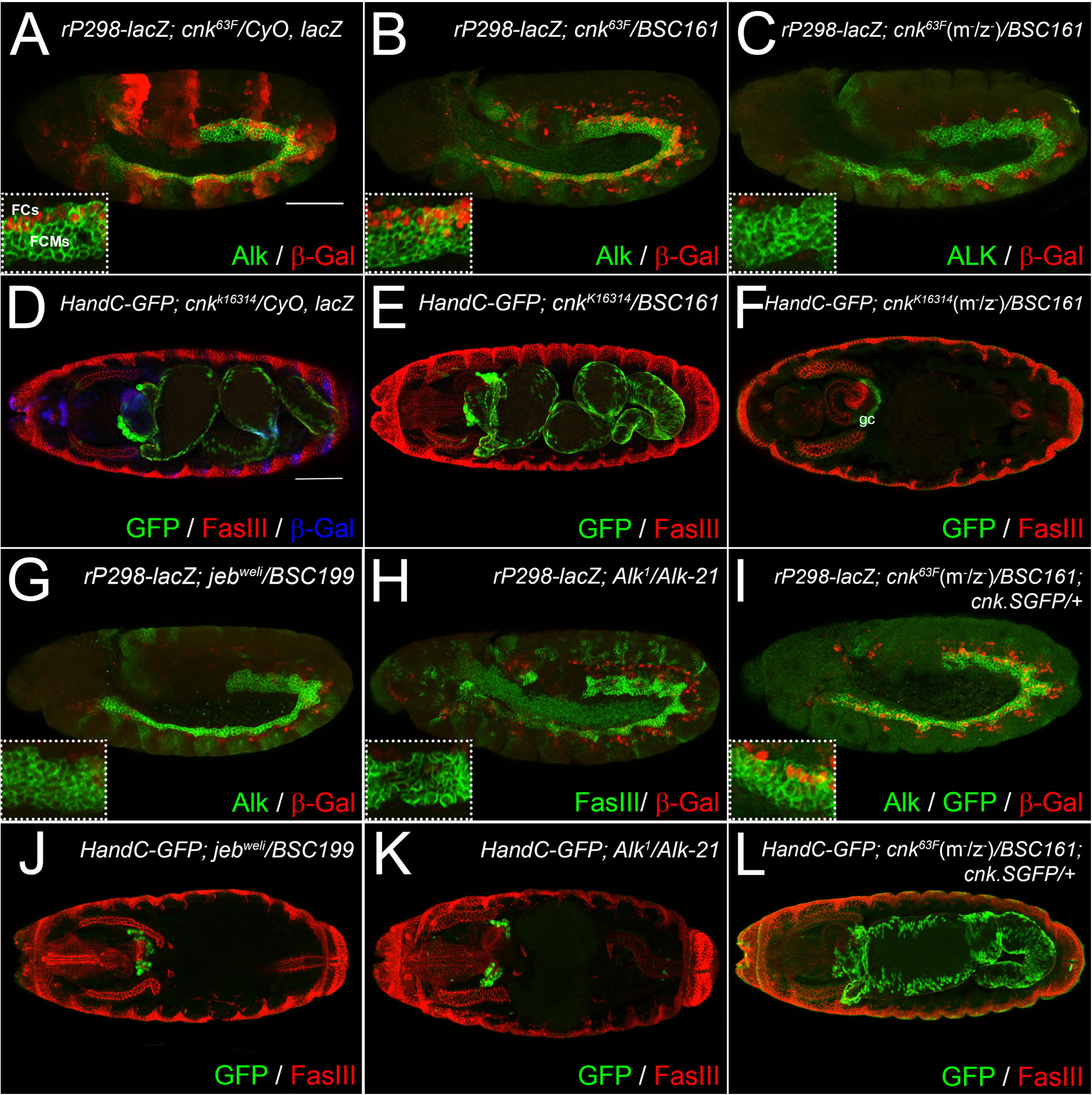
Loss of maternal and zygotic *cnk* genocopies *Alk* and *jeb* mutants. **(A-C, G, I)** VM of stage 11/12 *Drosophila* embryos expressing *rP298-lacZ* stained for β-Galactosidase (β-Gal in red) and Alk (green). In **(H)** the Alk staining is substituted by Fasciclin III (FasIII, green) staining. Control (A) and zygotic *cnk^63F^/BSCl6l* mutant embryos (B) exhibit a *rP298-lacZ-*positive FC row (FCs, FCMs are fusion competent myoblasts). In contrast, visceral FCs are absent in *cnk^63F^*(m^−^/z^−^)*/BSCl6l*embryos lacking both maternal and zygotic *cnk* (C) as well as in ***jeb*** (G) and Alk-mutant (H) embryos. FC formation in *cnk^63F^*(m^−^/z^−^)/BSCl6l can be restored by Cnk.SGFP (I, shown in green). **(D-F, J-L)** Stage 16 embryos carrying the *HandC-GFP* reporter stained for FasIII and GFP to reveal the presence of visceral muscles. A chambered gut surrounded by FasIII- and GFP-positive visceral muscles is present in control (D), zygotic *cnk^63F^/BSCl6l* (E) as well as “rescued” *cnk^63F^*(m^−^/z^−^)*/BSCl6l; Cnk.SGFP/+* (L, GFP in green) embryos. Visceral muscles are absent in *cnk^63F^*(m^−^/z^−^)*/BSCl6l* (F), *jeb-* (J) and Alk-mutant (K) embryos. GFP-expression in garland cells (gc) confirms the presence of the *HandC-GFP* reporter in F, J and K. Scale bar: 50 μm.

### The *semang* (*sag*) complementation group consists of novel *cnk* mutations

In earlier work, Zhang et al. (Zhang et al. 1999) conducted a modifier screen to identify suppressors of the *Src42A^Su(Raf)1^* allele that suppressed the lethality of the hypomorphic *Raf^C110^* (also referred to as *phf^C110^* and *Raf^1^*) mutation. Intriguingly, germ line clone-derived embryos of one of the identified suppressor mutations named *semang* (*sag*) displayed a midgut morphology reminiscent of the phenotypes observed in *jeb, Alk* and *cnk* (m^−^/z^−^) mutant embryos (Fig. 3; Fig. 3-figure supplement 1). We therefore analysed the *sag* mutations with respect to a possible role in visceral mesoderm formation. *sag* maps to 2R(54), a region that also harbours the *cnk* locus and, like *cnk, sag* is involved in the cell-autonomous specification of certain photoreceptor cells in the *Drosophila* eye, where it functions in the Ras/Raf/ERK pathway downstream of the EGF-receptor (Zhang and Lu 2000). Complementation tests revealed that the *semang* alleles *sag^13L^* and *sag^32-3^* failed to rescue the molecularly characterised mutation *cnk^k16314^*, as well as *Df(2R)BSC161* which deletes the entire *cnk* locus. Sequencing of the *cnk* locus in both *semang* alleles confirmed them to be *cnk* mutants at the molecular level (Fig. 3-figure supplement 1, see also Material and Methods). In *sag^13L^* mutants we detected a point-mutation (2R:C17415781T) that introduces a stop codon in place of Gln1156 of the Cnk protein while *sag^32-3^* mutants exhibit a nucleotide exchange (2R:T17415826A) leading to a premature stop in place of Lys1141. Notably, both stops are located between the RIR and the Src42A binding motif (pYELI) that has been suggested to relieve the inhibitory effect of Cnk on Raf-dependent MEK phosphorylation (Fig. 3-figure supplement 1) (Douziech et al. 2003) (Laberge et al. 2005). Therefore, the *sag* mutations are novel alleles of *cnk* and are henceforth referred to as *cnk^sag13L^* and *cnk^sag32-3^*.

### Cnk functions between Ras and Raf in the MAPK/ERK pathway

Given their possible role as active suppressors of Raf downstream of Alk we analysed germ line clone-derived, *cnk* (m^−^/z^−^) mutant embryos in the background of activated signalling pathway components. To do this, we employed *bap3-Gal4-driven* expression of the Alk ligand Jeb (*UAS-jeb*), an activated variant of the *Drosophila* Ras oncogene at 85D (*UAS-Ras85D^V12^*) and a constitutively active form of the *Drosophila* Raf kinase Polehole (*UAS-Raf.gof*) in the VM (Fig.4). As hypothesised and in agreement with previous studies (Lee et al. 2003) (Varshney and Palmer 2006) (Wolfstetter et al. 2009) (Popichenko et al. 2013), expression of these factors in the VM resulted in an increase in *HandC-GFP* expression and ERK phosphorylation (pERK) within the VM suggesting that the majority of Alk-positive visceral myoblasts were converted into FCs (Fig. 4C, E and G). Maternal and zygotic loss of Cnk function in *cnk^sag13L^*(m^−^/z^−^)*/BSC161* embryos was sufficient to abolish both ERK phosphorylation and *HandC-GFP* expression in the VM (Fig. 4B). In agreement with a role for Cnk downstream of the activated Alk receptor, *cnk^sag^* alleles supressed the over-expression effects of Jeb and *Ras^V12^* in *HandC-GFP;* cnk(m^−^/z^−^); *bap3>jeb* and *HandC-GFP; cnk*(m^−^/z^−^)*; bap3>Ras^V12^* embryos (Fig. 4D and F), but not that of activated Raf in *HandC-GFP; cnk*(m^−^/z^−^); *bap3>phl^gof^* embryos (measured as ERK phosphorylation or HandC-GFP; Fig. 4H). These data suggest that Cnk functions downstream of Jeb, Alk and Ras but upstream of Raf in the Alk-signalling pathway.

**Figure 4.**
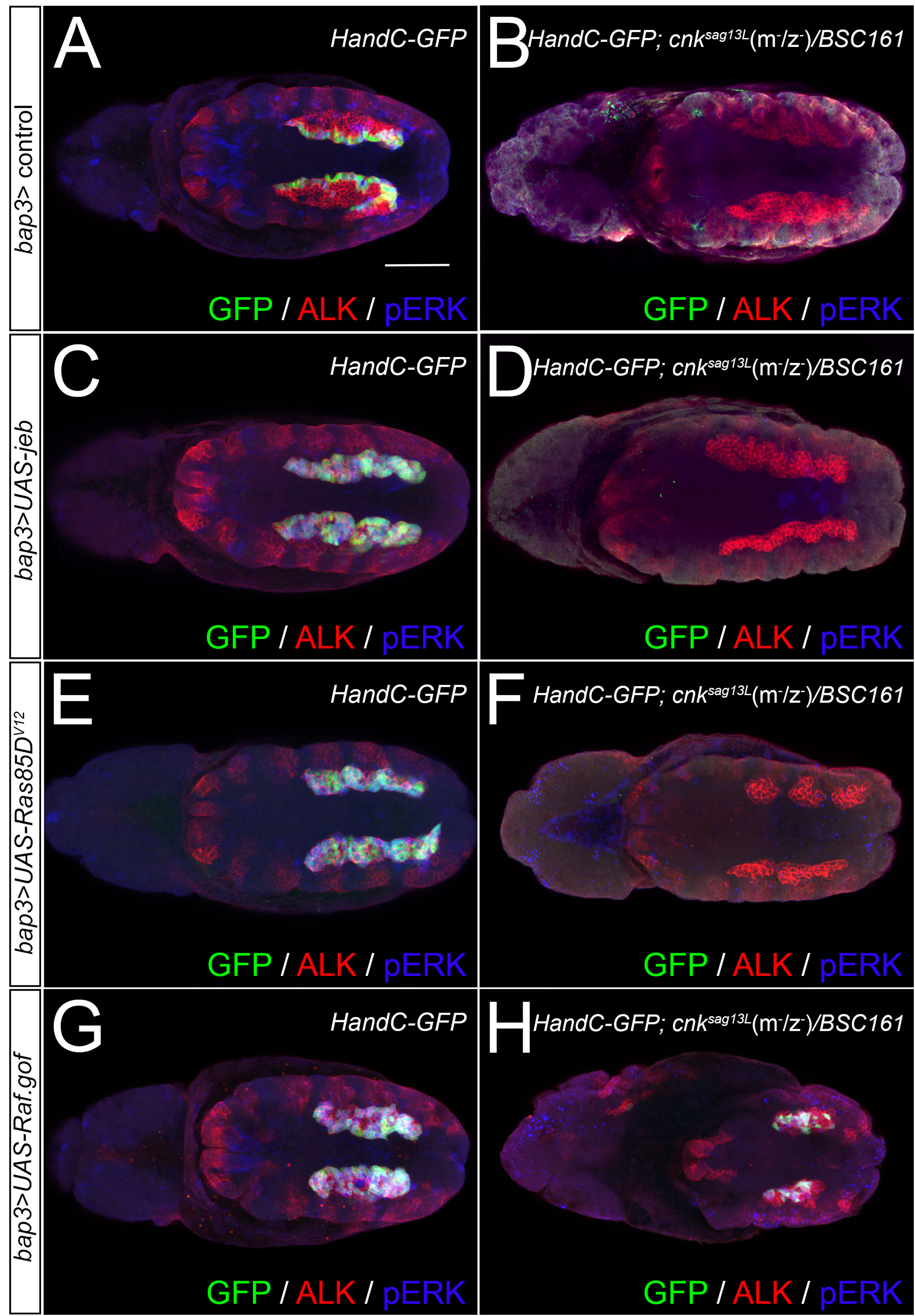
Cnk functions between Ras and Raf in the Ras/ERK pathway. Cnk is required for ERK activation downstream of Alk and Ras in the embryonic VM. Stage 11/12 embryos carrying the *HandC-GFP* reporter were stained for anti-pERK (shown in blue), GFP (green) and Alk (red) to evaluate Alk-signalling in the VM. Dorsal views are shown. **(A)** pERK staining and *HandC-GFP* reporter expression in the visceral FCs (arrow) of a control embryo. **(B)** Removal of maternal and zygotic *cnk* (cnk^sag13L^(m^−^/z^−^)/BSC161) results in a loss of both pERK and *HandC-GFP* expression. **(C, D)** pERK staining and *HandC-GFP* reporter expression is observed in all Alk positive VM cells of *bap3>UAS-jeb* embryos (C), but absent in *bap3>UAS-jeb* expressing embryos lacking maternal and zygotic *cnk (cnk^sag13L^*(m^−^/z^−^)*/BSCl6l).* (E) pERK staining and *HandC-GFP* reporter expression is observed in all Alk-positive VM-cells of *bap3>UAS-Ras85D^V12^* embryos. **(F)** Removal of maternal and zygotic *cnk (cnk^sag13L^*(m^−^/z^−^)*/BSCl6l*) results in a loss of both pERK and *HandC-GFP* expression in *bap3>UAS-Ras85D^V12^* expressing embryos. **(G)** pERK staining and *HandC-GFP* reporter expression is observed in the Alk-positive of *bap3>UAS-Raf.gof* embryos. **(H)** While less robust, both pERK staining and *HandC-GFP* reporter expression are observed in *bap3>UAS-Raf.gof* expressing maternal and zygotic *cnk (cnk^sag13L^*(m^−^/z^−^)*/BSCl6l*) mutants. Scale bar: 50 μm.

### The Cnk AIM-motif is required for robust activation of ERK in visceral FCs

The interaction between Alk and the c-terminal region of Cnk in the Y2H analyses defined a minimal AIM of 42 amino acids (Fig. 1A). Ectopic expression of Cnk^AIM^-tandem repeats resulted in the specific loss of visceral FC-identity, suggesting that this interaction is important for Alk-signalling. To test this hypothesis we employed CRISPR/Cas9 in combination with a *cnk^ΔAIM^*-donor construct to enforce homologous recombination with the *cnk* locus. This approach resulted in *cnk^ΔAIM^* mutants specifically lacking the 126 bp that encode for the minimal Cnk AIM (Fig. 5A, B). *cnk^ΔAIM^* mutants were viable and could be maintained as a fertile stock, suggesting that the minimal AIM is not a critical requirement for Alk-signalling in the VM. In agreement, VM-development in *cnk^ΔAIM^* mutants, examined with the FC-markers *rP298* or *HandC-GFP* as well as Alk or FasIII antibody staining, was indistinguishable from control embryos and did not reveal a loss of visceral FCs (Fig. 5B-F). We also employed antibody staining against pERK since activation of Alk by its ligand Jeb has been described to result in Ras/Raf/ERK pathway activation (Loren et al. 2001) (Lee et al. 2003) (Englund et al. 2003). Surprisingly, in contrast to the Alk-signalling output observed with the *rP298-lacZ* and *HandC-GFP* reporters, pERK levels were considerably decreased in the VM of *cnk^ΔAIM^* mutants when compared to wild-type controls (Fig 5F, G; quantified in I). Similar to *Alk^KO^* mutants (Fig 5H) ERK-activation was only affected in the visceral mesoderm (arrows in Fig 5F-H), while other tissues such as the tracheal pits (arrowheads in Fig 5F-H) still exhibited robust pERK staining. Thus, the VM-specific reduction of pERK indicated an important role for the Cnk AIM in Alk-dependent ERK-activation. The observation that visceral FC-specification proceeds in the context of reduced ERK-activation in the VM of *cnk^ΔAIM^* mutant embryos suggests the presence of (an) additional or extended binding interface(s) between Alk and Cnk or an additional indirect interaction with an unknown factor.

**Figure 5.**
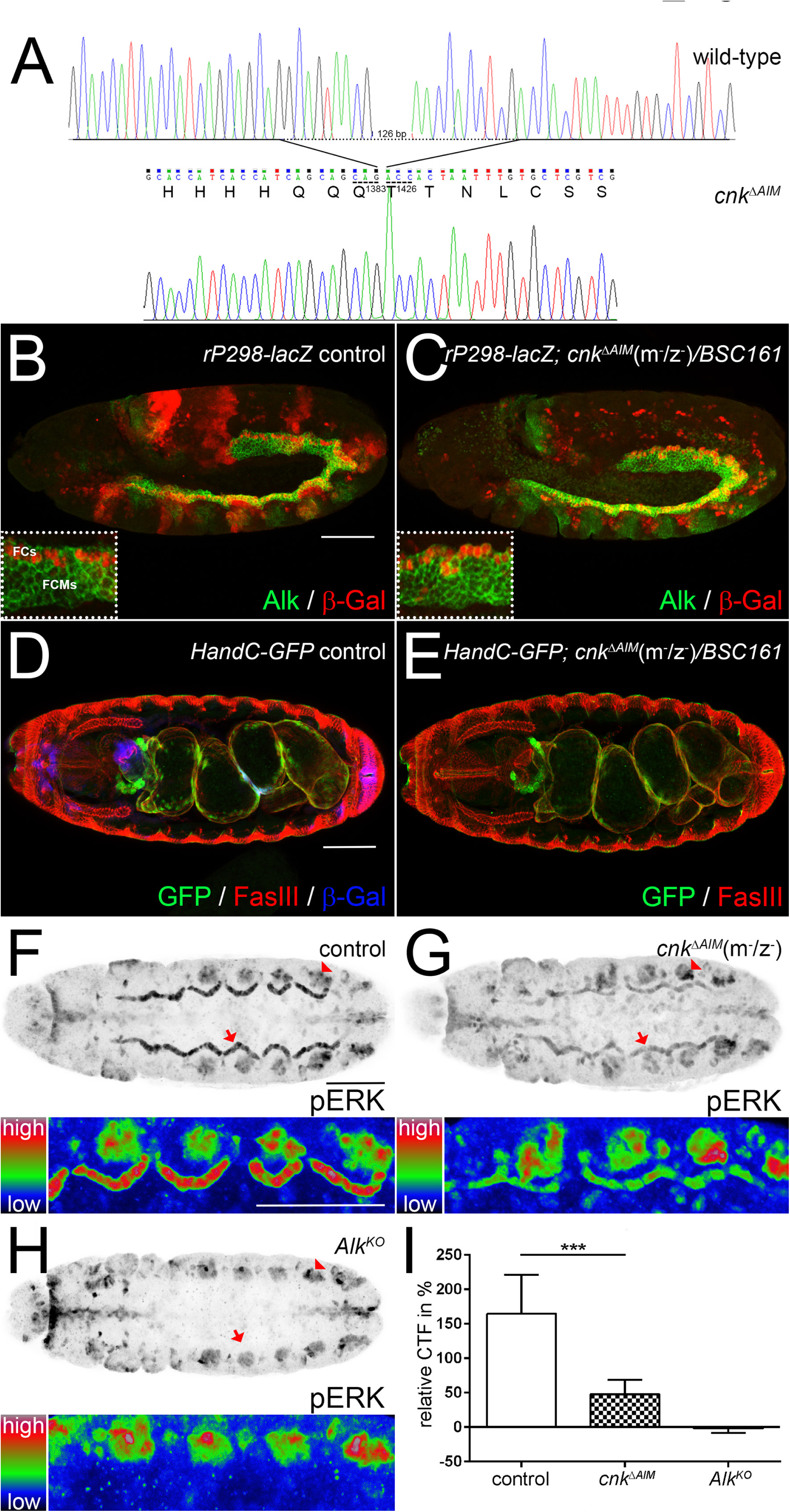
The Cnk AIM is required for robust activation of ERK in visceral FCs. (A) Chromatograms obtained by sequencing analysis of gDNA from *wild-type* and homozygous cnk^ΔAIM^ flies. The genome of cnk^ΔAIM^ flies harbours a 126 bp in-frame deletion in the endogenous *cnk* locus removing the region encoding for the 42 amino acid AIM between Gln(Q)1383 and Thr(T)1426 of the Cnk protein. **(B-E)** VM development proceeds normally in balanced siblings (B) and *cnk^ΔAIM^*(m^−^/z^−^)*/BSCl6l* embryos. Stage 12 embryos expressing the *rP298-LacZ* FC reporter stained for β-Gal (red) in FCs and Alk (green) in both visceral FCs and FCMs. **(D, E)** Stage 16 embryos stained for *HandC-GFP* reporter gene expression (green) and FasIII (red). β-Gal (blue) staining reveals balancer-associated LacZ-expression in a sibling embryo (D). **(F-H)** Phosphorylated ERK (pERK in black), detected by a phospho-specific MAPK antibody, in vFCs (arrows) and tracheal pits (arrowheads) of ventrally orientated stage 11 control (F), and *cnk^ΔAIM^* (m^−^/z^−^) (G) and *Alk^KO^* (H) embryos. The lower half of F, G and H depicts a (rainbow RGB colour LUT) heat map representation of close ups from the embryo in the upper half. pERK levels are specifically decreased in the VM of *cnk^ΔAIM^* (m^−^/z^−^) embryos (G) when compared to the control (F). No pERK staining is detectable in the VM of homozygous *Alk^KO^* (H) embryos. **(I)** Quantification of (background-)corrected total fluorescence intensities (CTF) in the VM relative to pERK-CTFs of the adjacent tracheal pits. Pairwise Student’s t-test was employed to reveal statistical significance (***=p<0,001, n=40 for each genotype). Scale bars: 50 μm.

### The small SAM-domain protein Aveugle is essential for Cnk activation downstream of Alk

The characterisation of an essential role for Cnk in Alk-mediated visceral FC-specification prompted us to search for and examine potential Cnk interacting partners. Therefore, we employed the Cnk full-length protein as bait and performed Y2H screening the same adult fly head cDNA prey library. Several Cnk binding partners were identified (Fig. 6A), including the small, sterile alpha motif (SAM)-domain containing protein Aveugle (Ave, also referred to as Hyphen or HYP) which has been shown to be important for MAPK/ERK-signalling (Douziech et al. 2006) (Roignant et al. 2006). Interestingly, we did not observe interactions with Raf, Ksr or Src42A which had previously been identified as binding partners of truncated Cnk variants (Therrien et al. 1998) (Laberge et al. 2005) (Douziech et al. 2003) (Douziech et al. 2006). To functionally analyse the role of the Cnk/Ave-interaction in more detail, we generated a series of sequence aberrations in the *ave* locus employing CRISPR/Cas9 genome editing. Two classes of molecular lesions were obtained: (i) deletions removing the ATG start codon of *ave* and (ii) smaller sequence aberrations that introduced shifts in the open reading frame (Fig. 6B). Since the *ave* locus partially overlaps with one isoform of the minus-orientated *Rpn6* gene, we confirmed that the generated *ave* alleles were able to complement the lethal, hypomorphic *Rpn6^20F^* mutation (Lier and Paululat 2002) as well as the lethal P-element insertion *Mi{ET1}Rpn6^MB09493^*. Homozygous as well as trans-heterozygous *ave* mutants survived until pupal stages suggesting that possible embryonic phenotypes are masked by a maternal component which had indeed been reported for *ave* (Roignant et al. 2006) (Graveley et al. 2011). We therefore analysed the visceral morphology of embryos lacking both maternal and zygotic Ave (*ave^CC9^*(m^−^/z^−^)) employing *HandC-GFP* as visceral FC-marker and antibody staining against FasIII to label the VM (Fig. 6F-H). In contrast to sibling embryos that had received one functional copy of *ave* and displayed wild-type VM morphology (Fig. 6C-E), *ave^CC9^*(m^−^/z^−^) embryos exhibited all hallmarks of the visceral phenotypes observed in *cnk*(m^−^/z^−^), *jeb* or *Alk* mutants such as loss of visceral FC identity (Fig. 6F), scattering of the visceral mesoderm after germ band retraction (Fig. 6G; arrowheads) and the absence of a functional gut musculature at late embryonic stages (Fig. 6H). Therefore we concluded that Ave could indeed serve as critical activator of Cnk in the Jeb/Alk-pathway.

**Figure 6.**
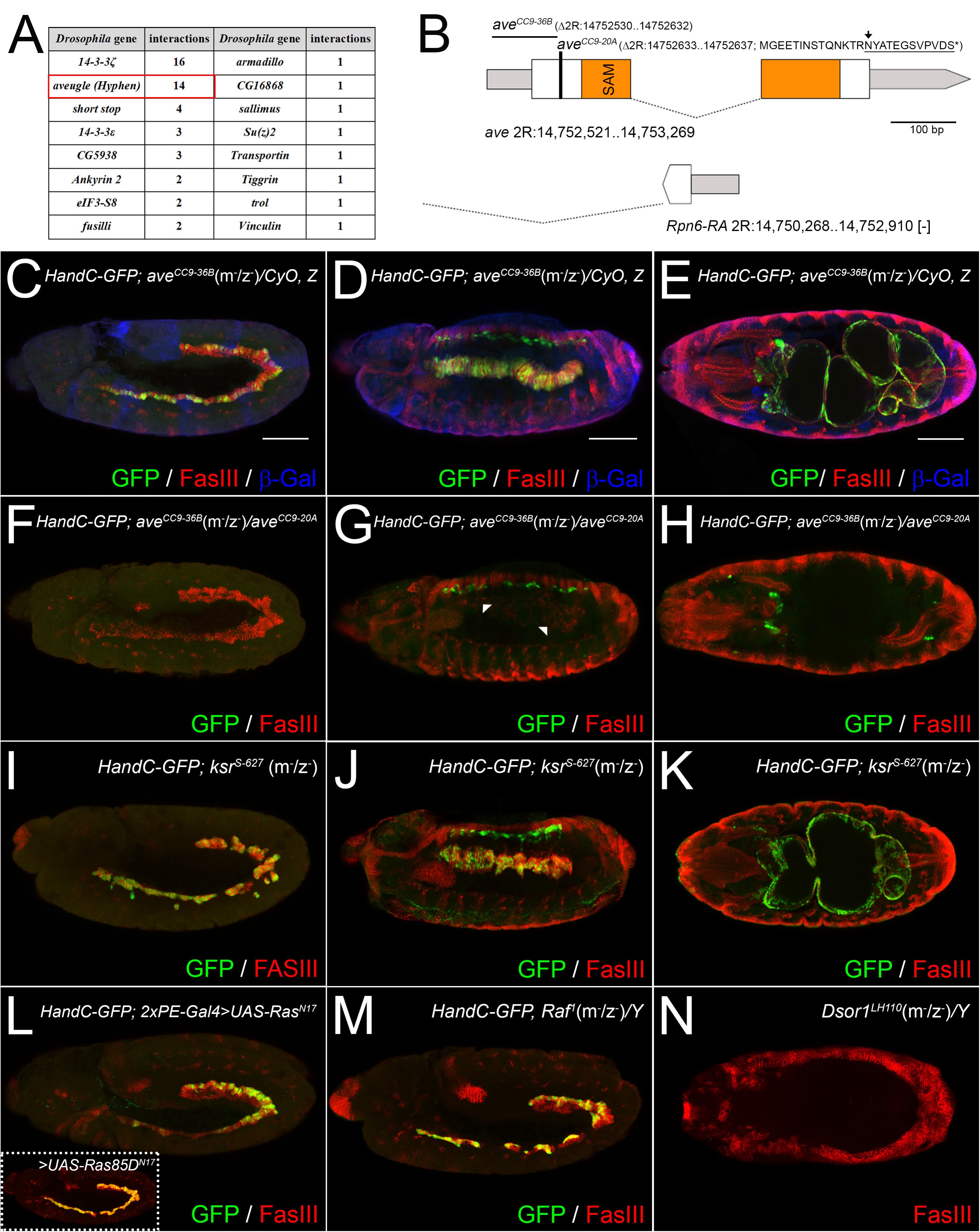
Ave, but not Ksr is required for Cnk-mediated Alk-signalling in the VM. **(A)** Table summarising binding partners and numbers of interacting preys obtained by Y2H screening employing full-length Cnk as bait. **(B)** Schematic representation of the *ave* locus. The first exon of the minus orientated Rpn6 gene is shown below; the CRISPR/Cas9-induced molecular lesions of *ave^CC9-20A^* and the *ave^CC9-36A^* null allele are highlighted above the *ave* locus. **(C-N)** Antibody staining against *HandC-GFP* reporter gene expression (green), FasIII (red) and balancer-associated β-Gal-expression (blue in C-E) in *Drosophila* embryos at stage 12 (C, F, I, L and M), 13/14 (D, G and J) and at the end of embryogenesis (E, H, K and N). Balanced *ave^CC9-36A^* sibling embryos specify visceral FCs (C), and VM development proceeds normally (D, E) while *ave^CC9-36A^(m/z˜)/ave^CC9-20A^* embryos exhibit loss of visceral FCs (F), disintegration of the VM (G, arrowheads) and absence of the visceral muscle (H). Visceral FCs are specified (I) in germ line clone derived *ksr^S-627^*(m^−^/z^−^) embryos that exhibit imperfect VM-stretching (J) and midgut constriction (K) at later stages. HandC-GFP-positive visceral FCs are present in embryos ectopically expressing mammalian or *Drosophila* (inset) Ras^N17^ under the control of the 2xPE-Gal4 driver (L) and in *Raf^1^* (m^−^/z^−^)*/Y* mutants (M). Embryos lacking maternal and zygotic (m^−^/z^−^) *Dsor1* (*Drosophila MEK*) exhibit the absence of differentiated internal structures as revealed by FasIII staining (N). Scale bars: 50 μm.

### Cnk function in the visceral mesoderm is independent of Ksr

To identify additional factors which are recruited by Ave-activated Cnk and might function in Jeb/Alk-signalling we investigated the role of Ksr. In agreement with previous findings, we did not observe obvious somatic clones of either *ksr^S-638^* or *ksr^S-627^* in germ line clone-producing females (Therrien et al. 1995) and only a small number of *ksr* (m^−^/z^−^) embryos could be obtained. Our analysis of *ksr^S-638^* and *ksr^S-627^* germ line clone-derived *ksr* (m^−^/z^−^) embryos carrying the *HandC-GFP* reporter did not reveal any loss of visceral FCs (Fig. 6I). In contrast to *cnk* (m^−^/z^−^) and *ave* (m^−^/z^−^) embryos, *ksr* (m^−^/z^−^) mutants developed visceral muscles although they exhibited milder phenotypes such as myotube stretching defects at stage 13/14 as well as an incomplete midgut constriction at the end of embryogenesis (Fig. 6J, K). To our surprise, terminal defects were present but variable and not as severe as previously reported for homozygous *huckebein (hkb)* and *tailless* (*tll*) embryos (Wolfstetter et al. 2009), suggesting that Torso-signalling and the expression of its downstream effectors, *hkb* and *tll*, is not entirely Ksr-dependent. Notably, terminal defects as well as the visceral muscle phenotype were more severe in *HandC-GFP*; *ksr^S-^*^*638*^(m^−^/z^−^)/*ksr^S-627^* embryos (Fig. 6-figure supplement 1) when compared to *HandC-GFP*; *ksr^S-^*^*627*^(m^−^/z^−^)/*ksr^S-627^.* Sequencing analysis revealed that *ksr^S-627^* is caused by a premature stop codon likely resulting in a truncated Ksr protein that lacks its entire kinase domain as well as the Raf-dimerization surface (Fig. 6-figure supplement 1) (Rajakulendran et al. 2009). Therefore it could be possible that the point mutations A696V and A703T identified in *ksr^S-638^* (Therrien et al. 1995) create a protein that dominantly interferes with Torso-signalling. Our analysis of embryos lacking Ksr, which exhibit HandC-GFP-positive FCs and the formation of a midgut, suggests that - in contrast to Cnk and Ave - Ksr is not a critical component of Jeb/Alk-signalling in the visceral mesoderm.

Ksr requires the presence of additional factors such as activated Ras to stimulate MEK-phosphorylation (Therrien et al. 1996) and ectopic expression of Ras^V12^ in the VM results in an excess of visceral FCs (Lee et al. 2003) (this study). In order to dissect the contribution of Ras to the pathway we ectopically expressed dominant inhibitory mammalian Ras^N17^ pan-mesodermally with *2xPE-Gal4* (Fig. 6L). Surprisingly, this did not result in a severe reduction of FCs although loss of FCs was observed upon ectopic expression of Cnk^AIM^ with the same driver line (Fig. 2G, H). Similar results were obtained upon ectopic expression of *Drosophila* Ras85D^N17^ (Fig. 6L, inset). To further address this finding, we employed the hypomorphic *Raf^1^* allele caused by an Arg>Leu amino acid substitution that corresponds to the well-known *Raf1^R89L^* mutation of human Raf1/c-Raf (Perrimon et al. 1985) (Fabian et al. 1994) (Hou et al. 1995). Comparable to Raf1^R89L^, the protein encoded by *Raf^1^* is unable to bind *Drosophila* Ras85D (Hou et al. 1995). In line with our previous experiments, we observed HandC-GFP-positive visceral FCs in germ line clone derived *Raf^1^*(m^−^/z^−^) embryos (Fig. 6M). On the other hand, germ line clone derived (m^−^/z^−^) embryos carrying the *Downstream of raf1* (*Dsor1*, encoding the single MEK of *Drosophila*) mutation *Dsorl^LHI10^* lacked most FasIII-positive internal structures such as VM, trachea, fore-and midgut and only rudiments of hindgut and salivary glands could be observed indicating that - in contrast to Cnk, Ave and Ksr - Dsor1/MEK functions more generally in embryonic RTK signalling. Taken together, our analysis reveals an essential function for Ave and Cnk, but not Ksr, as core components of Alk signalling in embryonic VM development (Fig. 7).

**Figure 7.**
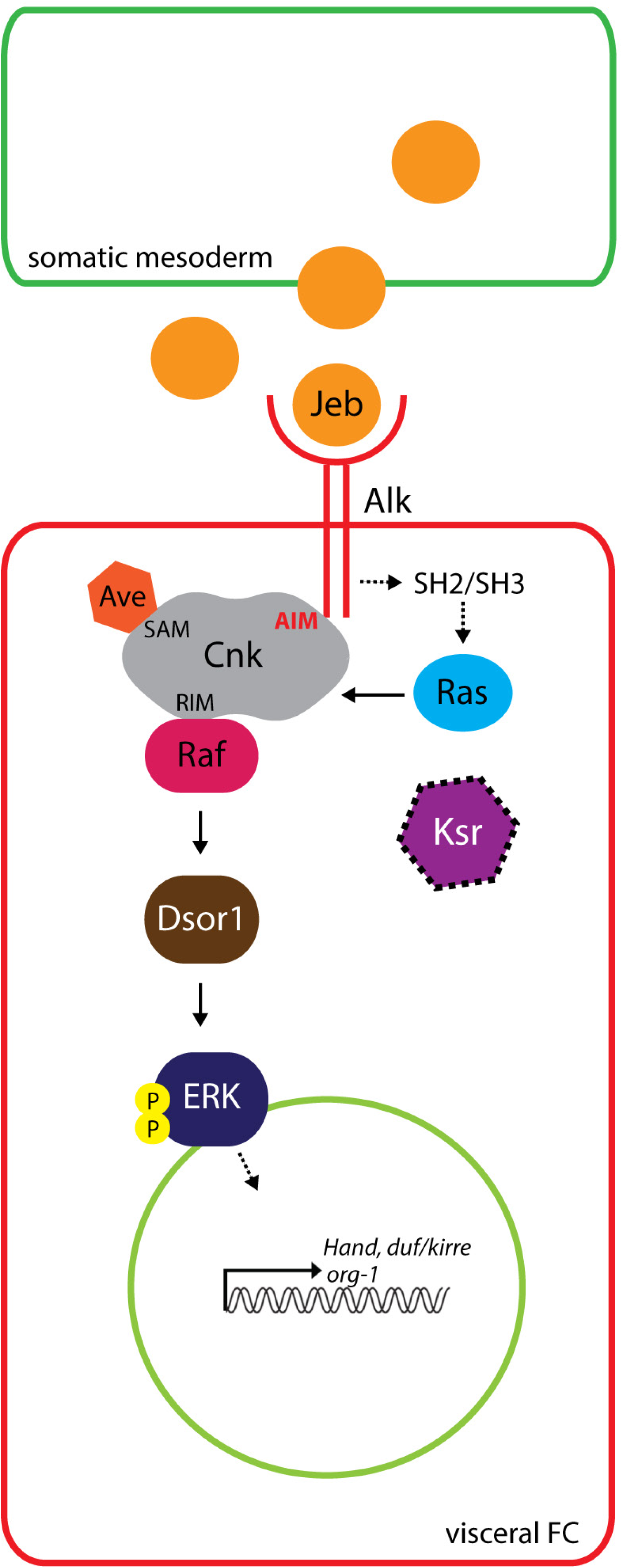
Current model for Alk-signalling during visceral FC-specification. Alk-activation at the membrane of a prospective visceral FC by the ligand Jeb, secreted from the neighbouring somatic mesoderm, induces the Ras/Raf/Mek/ERK signalling cascade, eventually leading to the transcriptional activation of downstream targets including *duf/kirre, org-1, Hand* among others. Cnk and Ave represent novel core components of the Alk-signalling pathway that are required downstream/in parallel to Ras and upstream of Raf to mediate visceral FC-specification. SH2/SH3 depicts a proposed adapter molecule that is likely to mediate Ras-activation upstream or in parallel to Cnk. Ksr is not essential for Jeb/Alk-signalling (indicated by dashed edges).

## DISCUSSION

In this study we have uncovered an essential function for the Cnk protein scaffold in Alk signalling. The identification of multiple Cnk-preys as Alk^ICD^ interactors in our Y2H-analysis allowed definition of a minimal Alk interaction motif (AIM) sufficient for the Cnk-Alk interaction. This interaction is supported by the loss of FC-specification in the VM of *cnk* (m^−^/z^−^) mutants, genocopying the embryonic *Alk* loss of function phenotype. The dominant negative effect of ectopic Cnk^AIM^-expression on Alk-signalling, together with the tissue-specific decrease of ERK-phosphorylation in the visceral FC row of *cnk^ΔAIM^* mutants strongly suggests that the function of Cnk in Alk-signalling is facilitated by a direct interaction between these two molecules. To our knowledge this is the first evidence for a direct interaction between Cnk and a RTK although various interactions between Cnk and membrane associated factors have been reported. In mammalian cell culture experiments, CNK1-promotes insulin signalling by binding to and regulating cytohesins (Lim et al. 2010) and binding of CNK1 to the transmembrane ligand EphrinB1 but not the EphB-receptor has been shown to link fibronectin-mediated cell adhesion to EphrinB-associated JNK signalling (Cho et al. 2014). Moreover, mammalian CNK2 (also reported as MAGUIN) which shares the greatest homology to *Drosophila* Cnk has been shown to bind various members of the membrane associated MAGUK-family proteins (Yao et al. 1999) as well as Densin-180, a LAP-family protein (Ohtakara et al. 2002). It will be interesting for future studies to determine whether the RTK binding capacities of Cnk are limited to Alk.

The Cnk protein localises in close proximity to the plasma membrane (Therrien et al. 1998) (Lanigan et al. 2003) (this study). While the Alk-Cnk^AIM^ interaction appears to be important for the robust activation of ERK downstream of Alk in the VM, Alk does not appear to be required for the subcellular localisation of Cnk. Whether the Alk-Cnk interaction is regulated by Alk activity, potentially resulting in post-translational modification of Cnk will be interesting to pursue in further studies. Notably, tyrosine phosphorylation of *Drosophila* Cnk has been reported upon co-expression with the activated Sevenless (SEV^S11^) RTK in S2 cells (Therrien et al. 1998) (Laberge et al. 2005) and activation of the Platelet-derived growth factor receptor (PDGFR) induces tyrosine phosphorylation of mammalian CNK1 leading to changes in CNK1 subcellular localization (Fischer et al. 2015).

Although Cnk is generally expressed in the *Drosophila* embryo, it appears to be of particular importance for Alk signalling (Fig. 7), perhaps facilitated by direct binding of Cnk^AIM^ to the Alk^ICD^. In agreement with earlier analysis of embryonic Torso signalling in *ave* (m^−^/z^−^) mutants (Roignant et al. 2006), we observed that terminal structures were formed in both *cnk* and *ave* mutants. Additional support for a differential importance for Cnk among RTKs is provided by the fact that Heartless RTK-mediated induction of the VM lineage (Beiman et al. 1996) (Gisselbrecht et al. 1996) (Shishido et al. 1997) is not conspicuously affected in either *cnk* or *ave* (m^−^/z^−^) mutant embryos. Moreover, selectivity for a requirement of CNK by different RTKs has also been observed in mammalian cells, as CNK2 appears to be required for NGF- but not EGF-induced ERK activation in PC12 cells (Bumeister et al. 2004). Therefore, while Cnk contributes to multiple signalling events it appears to be of preferential importance to different RTKs, perhaps reflecting differential wiring of downstream signalling in different developmental processes (Sopko and Perrimon 2013), an aspect that will be interesting to explore in future studies.

Cnk has been described as protein scaffold that facilitates RAS/RAF/MAPK signalling at the plasma membrane, allowing signal integration to enhance Raf and MAPK activation (Claperon and Therrien 2007). The epistatic analysis presented here shows that Cnk is required downstream of the activated Alk receptor and Ras^V12^, but upstream of activated Raf in the VM (Fig. 7), which is in agreement with previously reported studies in *Drosophila* (Therrien et al. 1998) (Walker et al. 2013). Thus, activated Ras seems to require Cnk to transmit signalling to Raf in the VM. Our observation that visceral FCs are specified upon ectopic expression of Ras^N17^ and also in *Raf^1^* (m^−^/z^−^) mutant embryos seemingly questions the importance of Ras in Alk-signalling in this context. The *Raf^1^* allele is caused by an amino acid substitution that resembles the mammalian *Raf^R89L^* mutation, and is unable to bind to Ras85D (Hou et al. 1995). However, the *Raf^R89L^* variant exhibits residual binding to CNK1 in human cells which is strongly enhanced in the presence of oncogenic Ras (Ziogas et al. 2005). Moreover, CNK has been reported to drive the compartmentalisation of Raf at the plasma membrane (Anselmo et al. 2002). Therefore, it is conceivable that direct binding to Ras might not be essential for Raf activation in the presence of a CNK scaffold. On the other hand, ectopic expression of Ras^N17^ interferes with “wild-type” Ras signalling at the guanine nucleotide exchange factor (GEF) level and requires an excess of the expressed transgene compared to the endogenous protein (Manser 2002). Unfortunately, our experiments do not allow us to quantify Ras^N17^ expression in the *Drosophila* VM. Additionally, the effects of Ras^N17^ might be masked by low Ras-GAP activities or the presence of multiple GEFs in the VM which could permit residual activation of endogenous Ras, eventually leading to FC-specification. There is also a possibility that Ras64B, the second Ras member in *Drosophila* or another GTPase like Rap1, could functionally substitute for Ras85D since Ras85D-independent Raf-activation has been observed for Torso signalling (Hou et al. 1995) (Raabe et al. 1996) (Mishra et al. 2005). In summary, additional experiments employing different sets of Ras and Raf mutations will be required to decipher their exact contribution to the Alk signalling pathway.

Aveugle/Hyphen (Ave/HYP) is a small, sterile alpha motif (SAM)-domain containing protein that functions in MAPK/ERK signalling between Ras and Raf (Douziech et al. 2006) (Roignant et al. 2006). Ave directly binds to Cnk via an interface formed by their SAM domains (Rajakulendran et al. 2008). Moreover, this interaction is thought to be necessary for the recruitment of Ksr to a complex which in turn promotes Raf activation in the presence of activated Ras (Douziech et al. 2006). While we have clearly identified Ave as critical component for Cnk function downstream of Alk, the single Ksr in *Drosophila* was not required for Alk-mediated FC-specification (Fig. 7). Ksr was initially identified in Ras-modifier screens and has been shown to stimulate MAPK/ERK activation in *Drosophila, C. elegans*, and different *in vitro* models (Kornfeld et al. 1995) (Sundaram and Han 1995) (Therrien et al. 1995) (Therrien et al. 1996) (Xing et al. 1997) (Roy et al. 2002). Furthermore, Cnk was originally identified in a Ksr-dependent genetic screen in *Drosophila* (Therrien et al. 1998) and its function has been proposed to mediate the association between Ksr and Raf (Douziech et al. 2006) suggesting that Ksr should also play an important role in Alk-signalling. However, the role of Ksr is unclear, with early reports suggesting an inhibitory function rather than an activating potential in RTK-signalling (Joneson et al. 1998) (Denouel-Galy et al. 1998). Ksr requires the presence of additional factors such as 14-3-3 proteins or activated Ras (Therrien et al. 1996) (Xing et al. 1997) and loss of Ksr-1 supresses Ras^E13^-induced but not wild-type signalling during *C. elegans* vulva formation suggesting altered affinities of Ksr for different variants of Ras (Sundaram and Han 1995). Therefore, we cannot exclude the possibility that –albeit non-essential-Ksr might enhance Alk-signalling by integrating signals from an activated Ras-variant to Cnk-associated Raf.

The importance of ERK activation in RTK-mediated signalling in the *Drosophila* embryo is difficult to address directly since the *rolled* locus (encoding the only MAPK/ERK1,2 orthologue in *Drosophila*) is located close to the centromere and therefore not accessible for standard germ line clone analysis. The removal of the AIM from Cnk by genomic editing of the *cnk* locus suggests that even reduced levels of pERK are sufficient *in vivo* to promote Alk-induced specification of visceral FCs. While it is clear that the AIM is sufficient to mediate the Alk-Cnk interaction, the ability of the *cnk^ΔAIM^* mutant to support Alk-driven FC-specification in the developing embryo highlights additional requirements *in vivo.* One possibility is that while the 42 amino acid AIM is sufficient to bind Alk, a larger region presents a more extensive binding interface *in vivo.* Another option is that additional interactions exist between Cnk, which is a large protein containing multiple domains, and Alk. Alternatively, other domains in Cnk may form indirect interactions with either additional Alk binding proteins and/or the PH domain of Cnk may provide additional interactions with the plasma membrane that contribute to the Alk-Cnk interaction *in vivo.*

In summary, we have identified a novel interaction between Alk and Cnk mediated by a minimal 42 amino acid motif that is required for robust activation of ERK by Alk in the VM of *Drosophila* embryos. Moreover, we show that Cnk is critically required for Alk-mediated specification of FCs in this tissue. Together with the small SAM-domain containing protein Ave/Hyp, Cnk represents an important signalling module that appears to be preferentially required for Alk-mediated signalling over other RTKs during embryogenesis. Indeed, Cnk and Ave represent the first molecules identified downstream of Alk whose loss genocopies the lack of visceral FC-specification of *Alk* and *jeb* mutants. Further work should allow a better understanding of the importance of Cnk in Alk-signalling and whether this is conserved to mammalian systems.

## MATERIALS AND METHODS

### *Drosophila* husbandry and fly crossings

Standard *Drosophila* husbandry procedures were followed (Ashburner 1989). Stocks were maintained on a potato-mash-based diet at room temperature. Crosses were performed at 25°C. LacZ- or GFP-balancer chromosomes (Bloomington *Drosophila* Stock Center at Indiana University, BDSC) were employed to distinguish the progenies of a cross. As “wild-type” controls we used *white^1118^* or balanced, heterozygous sibling embryos. Image stacks of adult flies were acquired with a Zeiss AxioZoom.V16 stereo zoom microscope equipped with LED ring light and an Axiocam 503 colour camera and further processed employing the enhanced depth of focus (EDF) module of the ZEN Blue edition software.

### Fly stocks

Fly stocks were obtained from BDSC (NIH P400D018537), the KYOTO Stock Center (DGRC) at Kyoto Institute of Technology and the Vienna *Drosophila* Resource Center (VDRC) at the Campus Science Support Facilities GmbH (CSF). Other lines used: *rP298-lacZ* (Nose et al. 1998), which is an enhancer trap in the *dumbfounded/kin of irre* locus (Ruiz-Gomez et al. 2000). *HandC-GFP* (Sellin et al. 2006), *bap3-Gal4* (Zaffran et al. 2001), *UAS-jeb* (Varshney and Palmer 2006), *UAS-Alk* (Loren et al. 2001), the EMS alleles *Alk^1^* and *Alk^10^* as well as the *Alk* deficiency *Df(2R)Alk-21* (Loren et al. 2003) and *jeb^weli^* (Stute et al. 2004) which carries a stop codon instead of Gln74 in the Jeb coding region (2R: C12117841T). In the *Alk^KO^* allele, exon 8 of the endogenous *Alk* locus has been replaced with an *attP* landing site following the protocol of Baena-Lopez et al. (Baena-Lopez et al. 2013). *Alk* exon 8 encodes for Asn1197-Cys1701 that correspond to the gross of the Alk^ICD^ (Tyr1128-Cys1701). *cnk^14C^* (also referred to as *cnk^C14^*), *cnk^63F^*, and *cnk^116C^* were identified in a mosaic genetic screen (Janody et al. 2004). Sequencing of these alleles revealed nucleotide exchanges in the coding region of *cnk* resulting in premature stops in place of Trp17 (2R:G17419418A in *cnk^63F^*), Gln253 (2R:C17418490T in *cnk^14C^*) and Trp806 (2R:G17416829A in *cnk^116C^).* The *semang* alleles *sag^323^* and *sag^13L^* (Zhang et al. 1999) were identified as novel *cnk* alleles (see results section for a detailed description) and are therefore referred to as *cnk^sag32-3^* and *cnk^sag13L^.* The *ave^CC9-20A^* allele carries a five nucleotide deletion (2R:C14752633-A14752637) that induces a frameshift resulting in a premature protein truncation (predicted sequence: MGEETINSTQNKTR<u>NYATEGSVPVDS*</u>, underlined amino acids differ from the wild-type Ave protein). In *ave^CC9-36B^*, 103 bp of the *ave* locus (2R:C14752530-A14752632) including the ATG start codon are deleted and eight nucleotides (AAACTACG) have been inserted between the Cas9 cutting sites. *ave^108V^* (Roignant et al. 2006) was used for complementation tests. The molecularly characterised allele *ksr^S-638^* (Therrien et al. 1995) and a so far uncharacterised allele, *ksr^S-627^* (Karim et al. 1996), were employed to generate germ line clone-derived embryos. Sequencing of *ksr^S-627^* revealed a nucleotide exchange (3R:C5483151T) indicating a pre-mature stop in place of Gln163 which results in a truncated protein lacking its kinase domain and Raf dimerization surface (Fig. 6-figure supplement 1).

### Generation of somatic clones and germline mosaics

Germ line clones were generated combining the FLIP-recombinase/FLP-recombination-target (FLP/FRT) system (Golic and Lindquist 1989) and the ‘dominant-female-sterile’ technique (DFS) as described in Chou et al. (Chou et al. 1993) and Chou and Perrimon (Chou and Perrimon 1996).

### Whole mount *in situ* hybridisation and fluorescent antibody staining

Embryo staining was carried out according to Müller (Müller 2008). For staining of phosphorylated MAPK/ERK (pERK) we followed the protocol from Gabay et al. (Gabay et al. 1997). Whole mount *in situ* hybridisation was done according to Lécuyer et al. (Lecuyer et al. 2008) with modifications adapted from Pfeifer et al. (Pfeifer et al. 2014). The following antibodies were used in the specified dilutions: guinea-pig anti-Alk (1:1000) (Loren et al. 2001), rabbit anti-Alk (1:1000) (Loren et al. 2001), chicken anti-β-Galactosidase (1:200; Abcam ab9361), rabbit anti-β-Galactosidase (pre-absorbed on fixed embryos, 1:1500; Cappel #0855976), rabbit anti-GFP (1:500; Abcam ab290), chicken anti-GFP (1:300; Abcam ab13970), mouse anti-activated MAPK/-diphosphorylated ERK1&2 (1:250; Sigma #M8159), and mouse 16B12 anti-HA.11 (1:500; Covance #MMS-101P). The monoclonal antibody 7G10 anti-Fasciclin III (1:50) was obtained from the Developmental Studies Hybridoma Bank (DSHB). Alexa Fluor®-, Cy^TM^-, Biotin-SP-, and HRP-coupled secondary antibodies as well as animal sera were purchased from Jackson ImmunoResearch. For signal amplification we used the Vectastain Elite ABC kit (Vector laboratories) in combination with Tyramide Signal Amplification (TSA^TM^) Plus Fluorescein or Cyanin3 systems (Perkin Elmer). Stained embryos were dehydrated in an ascending ethanol series before clearing and mounting in methyl salicylate. Samples were analysed under a Zeiss Axio Imager.Z2 microscope and images were acquired with a Zeiss LSM800 confocal microscope or an Axiocam 503 colour camera employing ZEN Blue edition software.

### DNA constructs

*UAS-5xcnk^AIM^.3xHA, UAS-Myr^src42A^⸬5xcnkAIM*, and *UAS-Myr^src42A^⸬5xcnk^AIM^.3xHA* DNA-sequences were assembled by GenScript and encode for five tandem repeats of the Alk interaction motif of Cnk (Cnk^AIM^; Ala1385-Ser1425). The myristoylation signal (Met1-Lys10) from *Drosophila* Src42A was employed to force membrane localisation of the respective constructs whereas a c-terminal, triple hemagglutinin (3xHA)-tag was used to detect construct expression. The assembled sequence was cloned into the *pUAST* transformation vector (Brand and Perrimon 1993) employing *EcoRI* and *XbaI* restriction sites. To generate the *cnk^ΔAIM^* gene editing donor construct we PCR-amplified ~1 kb homology arms from genomic DNA of *w*; P{FRT(w^hs^)}G13* flies employing the following primer combinations: CCAGAGTCCCAGTAGCAAGTCGAGT and <u>AATTAGTGGTCTGCT</u>GCTGATGGTGATGG as well as <u>AGCAGACCACTAATT</u>TGTGCTCG in combination with TTAGGTCTTTGAATAAGTTGCGTGC. This introduced a 15 bp overlapping sequence (underlined in the primer sequences) which was further used as “internal primer” in an assembly-PCR reaction with the external (non-underlined) primer pair. The obtained *cnk^ΔAIM^* donor sequence was sub-cloned in *pCRII* (Invitrogen) and shuttled into *pBluescript II KS(-)* employing *HindIII* and *XhoI* restriction sites. All DNA-constructs were verified by Sanger sequencing (GATC Biotech).

### Generation of *cnk^ΔAIM^* mutants employing CRTSPR/Cas9-facilitated homologous recombination

To induce double strand breaks in the region of the Cnk/Alk interacting motif (Cnk^AIM^), we employed the sgRNA target sequence GCACAAATTAGTGGTCGAGG<u>TGG</u> (the Proto-spacer Adjacent Motif or PAM is underlined) which was cloned into the *pBFv-U6.2 gRNA* expression vector (GEPC). To achieve molecularly defined *cnk^AIM^* deletions (*cnk^ΔAIM^*)we forced homologous recombination by simultaneously injecting (BestGene Inc.) the sgRNA and the *pBluescript-cnk^ΔAIM^* gene editing donor construct into embryos carrying the *P{FRT(w^hs^)}G13* insertion and expressing Cas9 in their germ line (*M{vas-Cas9}ZH-2A*). Balanced stocks were established from single flies carrying the FRT insertion and a possible recombination event and further analysed by PCR for the desired modification in the *cnk* locus.

### Generation of *ave* mutants by CRISPR/Cas9-mediated genome editing

To induce CRISPR/Cas9-mediated lesions in the *ave* locus, the following sgRNA sequences were used: CGGTCGCGTAGTTTTCGTTC<u>TGG</u> and AGCAACAAAACAAATAGTGA<u>TGG</u> (the PAM motif is underlined). *pBFv-U6.2* expression vectors containing these sequences (GEPC) were injected into *M{vas-Cas9}ZH-2A/+(or Y); P{FRT(w^hs^)}G13/+* embryos by BestGene. Balanced stocks were established from single flies carrying the FRT insertion and further screened by PCR for molecular lesions in the *ave* locus.

### Databases & bioinformatics

Information about *Drosophila* genetics is available on Flybase (http://flybase.org/), the Database of *Drosophila* Genes & Genomes (dos Santos et al. 2015). Genomic coordinates refer to the Dmel_Release_6 sequence assembly (Hoskins et al. 2015). The Fancy Gene v1.4 application (Rambaldi and Ciccarelli 2009) and the MyDomains image creator (Hulo et al. 2008) were used to create scale models of gene loci and proteins. Fluorescence intensity measurements were acquired with the Fiji distribution of ImageJ (Schindelin et al. 2012). In brief: Area, mean fluorescence and integrated density values were acquired from regions of interest (ROI) selected in confocal stacks of stage 11-12 embryos. The ROIs corresponded to a non-stained area (background), an arch of the VM (revealed by Alk staining in the *Alk^KO^* mutant) and the adjacent tracheal pit (tp) for each measurement. Three to four measurements were taken from each analysed embryo. The (background-)corrected total fluorescence (CTF = Integrated density ROI – (area ROI x mean fluorescence background)) of the VM-arch relative to the adjacent tp was calculated to minimize staining- or stage-dependent fluctuations. For statistical analysis we employed the GraphPad Prism 6 software.

### Yeast two-hybrid experimental analysis

Yeast two-hybrid EST-library screening was conducted by Hybrigenics. The Matchmaker Gold Yeast Two-Hybrid System (Clontech) was employed to validate the interactions of the *Drosophila* Alk pB66-bait (Tyr1128-Cys1701, n-terminally fused to the Gal4 DNA binding domain) with interacting *Drosophila* Cnk *pB6*-prey clones (encoding for Gly1303-Ser1425 or Gln1383-Asn1538 respectively, provided by Hybrigenics) and to determine the minimal <u>A</u>lk <u>i</u>nteracting <u>m</u>otif (AIM). Standard cloning techniques were employed to generate the *pP6* prey-Cnk^AIM^ constructs.

## AUTHOR CONTRIBUTIONS

GW and RHP conceived the project, designed the experiments and wrote the first draft of the manuscript. GW, KP, JRvD, FH and XL conducted the experiments. GW, KP, XL and RHP analysed the data. All authors contributed to the final version of the manuscript.

## ACKNOWLEDGEMENTS

We are indebted to Manfred Frasch, Anne Holz, Maria Leptin, Akinao Nose, Achim Paululat, Marc Therrien and Jessica Treisman for generously sharing fly stocks. We would like to thank the *Drosophila* Genomics Resource Center (DGRC, supported by NIH grant 2P40OD010949-10A1), the Genome Engineering Production Group (GEPG) at Harvard medical school and Hybrigenics for providing cDNAs, vectors and DNA-constructs as well as BestGene Inc. for excellent fly injection service. We are thankful for reagents supplied by the Developmental Studies Hybridoma Bank, created by the NICHD of the NIH and maintained at the University of Iowa, the Bloomington *Drosophila* Stock Center (NIH P40OD018537), the KYOTO Stock Center (DGRC) at the Kyoto Institute of Technology as well as the Vienna *Drosophila* Resource Center (VDRC) at the Campus Science Support Facilities GmbH (CSF). We are much obliged to our colleagues Anne Holz and Gautam Kao for their helpful suggestions and critical comments on the manuscript. This work has been supported by grants from the Swedish Cancer Society (2015/391 to RHP), the Children’s Cancer Foundation (15-0096 to RHP), the Swedish Research Council (2015-04466 to RHP), the SSF Programme Grant (RB13-0204 to RHP), the Göran Gustafsson Foundation (RHP2016), the American Cancer Society (#RPG-96-13504-DDC to XL), and partially by NIH (1 P50 DK57301-01 to XL). GW has been supported by postdoctoral fellowships from the Swedish Childhood Cancer Foundation (NC2014-0045) and the Sven & Lily Lawski Foundation. KP has been supported by a postdoctoral stipend from the Carl Tryggers Foundation (CTS KF15:1 5).

## Figure 1-figure supplement 1

**(A-C)** Eyes of control (A), homozygous *cnk^63F^* (B), and transheterozygous *cnk^sag13L^/cnk^sag32-3^* (C) female flies carrying the *fTRG1248* insertion (Cnk.SGFP). Cnk.GFP is able to rescue lethality caused by *cnk* mutations, however, flies exhibit a weak rough eye phenotype which is slightly stronger in the background of the proposed Raf-repressor alleles *cnk^sag13L^/ cnk^sag32-3^.* Scale bar: 50 μm.

Figure 3-figure supplement 1. (A) Schematic representation of the *cnk* locus (adapted and modified from (Claperon and Therrien 2007) including genome coordinates and depicting regions encoding for Cnk domains in different colours (SAM=sterile alpha motif in orange, CRIC=conserved region in Cnk in light green, PDZ=post synaptic density protein (PSD95), *Drosophila* discs large tumor suppressor (Dlg1), and zonula occludens-1 protein (zo-1) in magenta, PH=pleckstrin homology domain in blue, RIM=Raf interacting motif in yellow, IS=inhibitory sequence in brown, pYELI=phosphorylation site for Src42A in dark green, Alk interacting motif (AIM) in red). UTRs appear narrower and in grey, introns as dashed lines. The position of the molecular lesions identified in different *cnk* alleles and the predicted changes in the resulting Cnk proteins are indicated. **(B-F)** Absence of the visceral muscle in germ line clone derived late stage embryos carrying the HandC-GFP reporter and different *cnk* mutations *“in trans”* to the *cnk* deficiency *BSC161.* The embryos were stained for GFP reporter gene expression (green) and FasIII (red). Scale bar: 50 μm.

Figure 6-figure supplement 1. **(A)** Schematic representation of Ksr after (Therrien et al. 1995) and (Rajakulendran et al. 2009). The position of the proposed Raf/Ksr dimerization surface is indicated by a green line, position of the point mutations in *ksr^S-627^* and *ksr^S-638^* alleles and the predicted consequences for the Ksr protein are highlighted. CA=conserved area, KD=Protein kinase domain. **(B)** Antibody staining against GFP (green) and FasIII (red) reveals impaired VM stretching and terminal defects in a late stage *HandC-GFP; ksr^S-638^(m^−^/z^−^)/ksr^S-627^.* Scale bars: 50 μm.

